# Neural specialization for ‘visual’ concepts emerges in the absence of vision

**DOI:** 10.1101/2023.08.23.552701

**Authors:** Miriam Hauptman, Giulia Elli, Rashi Pant, Marina Bedny

## Abstract

Vision provides a key source of information about many concepts, including ‘living things’ (e.g., *tiger*) and visual events (e.g., *sparkle*). According to a prominent theoretical framework, neural specialization for different conceptual categories is shaped by sensory features, e.g., living things are neurally dissociable from navigable places because living things concepts depend more on visual features. We tested this framework by comparing the neural basis of ‘visual’ concepts across sighted (n=22) and congenitally blind (n=21) adults. Participants judged the similarity of words varying in their reliance on vision while undergoing fMRI. We compared neural responses to living things nouns (birds, mammals) and place nouns (natural, manmade). In addition, we compared visual event verbs (e.g., ‘sparkle’) to non-visual events (sound emission, hand motion, mouth motion). People born blind exhibited distinctive univariate and multivariate responses to living things in a temporo-parietal semantic network activated by nouns, including the precuneus (PC). To our knowledge, this is the first demonstration that neural selectivity for living things does not require vision. We additionally observed preserved neural signatures of ‘visual’ light events in the left middle temporal gyrus (LMTG+). Across a wide range of semantic types, neural representations of sensory concepts develop independent of sensory experience.

**Significance Statement:** Vision offers a key source of information about major conceptual categories, including animals and light emission events. Comparing neural signatures of concepts in congenitally blind and sighted people tests the contribution of visual experience to conceptual representation. Sighted and congenitally blind participants heard ‘visual’ nouns (e.g., ‘tiger’) and verbs (e.g., ‘sparkle’), as well as less visual nouns (e.g., ‘barn’) and verbs (e.g., ‘squeak’) while undergoing fMRI. Contrary to previous claims, both univariate and multivariate approaches reveal similar representations of animals and light emission verbs across groups. Across a broad range of semantic types, ‘visual’ concepts develop independent of visual experience. These results challenge theories that emphasize the role of sensory information in conceptual representation.

## Introduction

Studies of ‘visual’ concepts in people born blind offer a strong test of the contribution of sensory experience to concepts and their neural instantiation. Vision is an important source of information about multiple conceptual categories, including living things (e.g., animals, plants), visual events (e.g., ‘sparkle’, ‘glow’), and colors (Locke, 1690; Barsalou, 1999; Gallese & Lakoff, 2005; Richter & Zwaan, 2009; Humphreys & Forde, 2001; Striem-Amit et al., 2018; Bi, 2021). The current study asks whether neural specialization for two ‘visual’ categories, living things and light events, requires visual experience. These ‘visual’ categories span a wide range of semantic types—from concrete entities to events—and thus offer a broad perspective on the contribution of vision to the neural instantiation of concepts.

In the case of living things, watching animals such as elephants and blue jays offers information about their shape, color, texture, and behavior. Inspired in part by such observations, classic neuropsychological theories attributed semantic deficits for living things to damaged visual knowledge (Allport, 1985; Warrington & Shallice, 1984; Farah & McClelland, 1991; Gaffan & Heywood, 1993; Moss et al., 1997; Tranel et al., 1997; Martin et al., 2000; cf. Caramazza & Shelton, 1998; Caramazza & Mahon, 2003). According to such theories, visual features play a privileged role in concepts of living things, particularly animals and plants. As a result, damage to visual knowledge disproportionately impairs the living things category (Warrington & McCarthy, 1987; Humphreys & Forde, 2001).

Consistent with the idea that vision shapes living things concepts, knowledge of plant and animal appearance differs across sighted and congenitally blind people. Semantic similarity judgments about plants (fruits and vegetables) are influenced by color in sighted but not blind participants, whereas tool concepts are similar across groups (Connolly et al., 2007). Likewise, congenitally blind people show low agreement about animal colors and rate large animals (e.g., ‘bear’, ‘rhinoceros’) as less familiar than sighted people do (Kim et al., 2019). These data raise the question of whether neural specialization for living things depends on vision.

Most prior work investigating neural specialization for living things in blind individuals has focused on lateral ventral occipito-temporal cortex (VOTC), where sighted people show a preference for living things, including images of animals and humans (Kanwisher et al., 1997; Chao et al., 1999; O’Craven & Kanwisher, 2000; Grill-Spector et al., 2004; Konkle & Carmazza, 2013). In support of the idea that visual experience is necessary for living things specialization, VOTC selectivity for animals is either absent or weaker in blind relative to sighted people (Mahon et al., 2009; He et al., 2013; Mattioni et al., 2020; see Bi et al., 2016 for a review).

More recently, studies using tactile human faces and human vocalizations as stimuli have reported lateral VOTC responses in blind people (Pietrini et al., 2004; van den Hurk et al., 2017; Ratan Murty et al., 2020; Mattioni et al., 2020; Bola et al., 2022). However, it is unclear whether these responses are driven by living things (i.e., animacy) per se or by the distinctive communicative relevance of human faces and vocalizations. The latter is especially plausible given that visual cortices of people born blind respond selectively to language (e.g., Röder et al., 2002; Bedny et al., 2011; Collignon et al., 2013; Tian et al., 2023).

Recent studies with sighted people have identified responses to living things (people and animals) outside the VOTC in temporo-parietal cortex, particularly in precuneus (PC) (Fairhall & Caramazza, 2013a; 2013b; Fairhall et al., 2014; Deen & Freiwald, 2022; Peer et al., 2015; Elli et al., 2019). Unlike in VOTC, responses to living things in PC in sighted people are amodal: both words and images elicit similar responses (Fairhall & Caramazza, 2013a; Fairhall et al., 2014). In one experiment, a classifier trained on patterns of neural responses to images of faces generalized its performance to words describing people in PC, suggesting similar underlying representations (Fairhall & Caramazza, 2013a). Whether neural specialization for living things in temporo-parietal cortex, particularly PC, requires vision to emerge is unclear. One previous study with blind individuals reported greater activity for famous people compared to face parts in PC but did not establish a PC preference for people compared to nonliving entities (Wang et al., 2016). The present study investigates PC selectivity for living things by comparing neural responses to living (animal) and nonliving (place) nouns in congenitally blind and sighted adults. By comparing animal nouns rather than communication-oriented human stimuli (e.g., vocalizations, faces) to places, we separate animacy or ‘living things’ representations from communicative relevance.

The second ‘visual’ category examined in the current study is light events (e.g., ‘sparkle’). Unlike living things, light events can only be directly perceived through vision. Furthermore, unlike animals, which are concrete entities situated in space and encoded in most languages by nouns, light emission events are situated in time and encoded in most languages by verbs (Talmy, 1975; Langacker, 1987; Frawley, 1992). We can therefore ask whether blindness-related change and/or preservation generalizes across semantic types. Behaviorally, blind adults distinguish light events such as flash vs. glow based on intensity and periodicity, similar to sighted people (Landau & Gleitman, 1985; Lenci et al., 2013; Bedny et al., 2019). Whether neural representations of visual events are similar in blind and sighted people has not been tested.

Our primary hypotheses concerning light events focus on the left middle temporal gyrus and neighboring temporal areas (LMTG+), since these play an important role in event representation. The LMTG+ responds more to event verbs (e.g., ‘run’) and event nouns (e.g., ‘hurricane’) than to nouns that refer to entities (e.g., ‘strawberry’) (Bedny et al., 2014; Lapinskaya et al., 2016; Martin et al., 1995; Kable et al., 2002; 2005; Bedny & Thompson-Schill, 2006; Elli et al., 2019; Damasio & Tranel, 1993; Davis et al., 2004). Neighboring temporal regions also encode semantics of visual events (e.g., Isik et al., 2017; Wurm & Caramazza, 2019). Blind and sighted people show similar neural responses to motion event verbs (e.g., ‘run’ and ‘kick’) in LMTG+ (Noppeney et al., 2003; Bedny et al., 2012). However, unlike light emission events, motion events can be experienced through sensory modalities outside of vision (e.g., audition, somatosensation) (Kemmerer et al., 2008; Pulvermüller & Fadiga, 2010; Yee, Chrysikou, & Thompson-Schill, 2017). Here we test whether LMTG+ responses to purely visual events (i.e., light emission verbs) develop differently in the absence of vision.

Three prior studies examined the neural basis of ‘visual’ concepts in blindness using color adjectives (e.g., ‘blue’), and one also used weather words like ‘rainbow’ (Striem-Amit et al., 2018; Wang et al., 2020; Bottini et al., 2020). These studies found evidence for blindness-related changes in VOTC and partial preservation in the anterior temporal lobe (ATL). However, the degree to which these findings apply broadly across ‘visual’ concepts remains unclear. Unlike weather and color words, events and living things elicit robust and distinctive neural signatures in sighted people, making it possible to test whether these signatures depend on vision (e.g., Fairhall & Caramazza, 2013a; Deen & Freiwald, 2022; Bedny et al., 2014; Elli et al., 2019).

The present study uses individual-subject univariate and multivariate approaches to compare neural representations of ‘visual’ categories across sighted and congenitally blind people. We first localized brain regions that exhibited a preference for entities vs. events in individual participants. Within entity- and event-preferring networks, we compared ‘visual’ to non-visual categories, i.e., conceptual types from the same class (entity or event) for which vision is thought to play a less important role. Congenitally blind and sighted participants performed within-category semantic similarity judgments on words from four entity/noun categories (living: birds, mammals; nonliving: manmade places, natural places) and four event/verb categories (visual: light, nonvisual: sound, hand action, mouth action). Using place nouns as our nonliving things category additionally enabled us to test for previously documented place selectivity in the medial VOTC of blind and sighted people (i.e., parahippocampal place area (PPA); Epstein & Kanwisher, 1998; Wolbers et al., 2011; He et al., 2013; van den Hurk et al., 2017; Mattioni et al., 2020).

A secondary goal of the current study was to examine responses to spoken words in the visual cortices of sighted and congenitally blind adults. Prior studies find responses to sentences and words in occipital cortices of congenitally blind people and to some degree even sighted people (Röder et al., 2002; Amedi et al., 2003; Bedny et al., 2011; Lane et al., 2015; Kanjlia et al., 2016; Bola et al., 2017; Seydell-Greenwald et al., 2020). We used univariate and multivariate methods to test the sensitivity of visual cortices to word meanings in both populations.

## Methods

### Participants

Twenty-one congenitally blind adults (13 females, age range 18-67 years, M = 39.14 ± 13.81 SD) and twenty-two sighted age and education matched controls (16 females, age range: 19-62 years, M = 37.55 ± 13.25 SD) participated in the experiment (Supplementary Table 1). Blind participants lost their sight due to pathologies of the eyes or optic nerve, anterior to the optic chiasm (i.e., not due to brain damage), and had at most minimal light perception since birth. Throughout the experiment, all participants (sighted and blind) wore a light exclusion blindfold to match their visual input. Sighted and blind participants were screened for cognitive and neurological disabilities (self-report). Participants gave written informed consent and were compensated $30 per hour. The study was reviewed and approved by the Johns Hopkins Medicine Institutional Review Boards. Four additional blind participants were scanned but excluded from the final sample because they were older than 70 years of age (n=2), they were not blind since birth (n=1), or they gave similarity judgments different from those of the group (n=1, correlation with the group lower than 2.5 SDs from the average for both verbs and nouns).

### Stimuli and procedure

While undergoing functional magnetic resonance imaging (fMRI), participants heard pairs of words and judged how similar the two words were in meaning on a scale from 1 (not at all similar) to 4 (very similar), indicating their responses via button press. Word stimuli consisted of 18 words in each of 8 semantic categories (Figure 1, Supplementary Table 2, see Supplementary Materials for full list of stimuli): 4 categories of entities/nouns (birds, e.g., ‘the crow’; mammals, e.g., ‘the fox’; manmade places, e.g., ‘the barn’; natural places, e.g., ‘the swamp’), and 4 categories of events/verbs (light emission, e.g., ‘to sparkle’; sound emission, e.g., ‘to squeak’; hand-related actions, e.g., ‘to pluck’, mouth-related actions, e.g., ‘to bite’). Words were matched across several linguistic variables (see Elli et al., 2019, and Supplementary Materials for details). Word pairs were presented in blocks of 4 and were grouped by semantic category within blocks. Each word appeared once within a block. Blocks were 16 s long and were separated by 10 s of rest. The experiment included a total of 144 blocks evenly divided into 8 runs.

**Figure 1:**
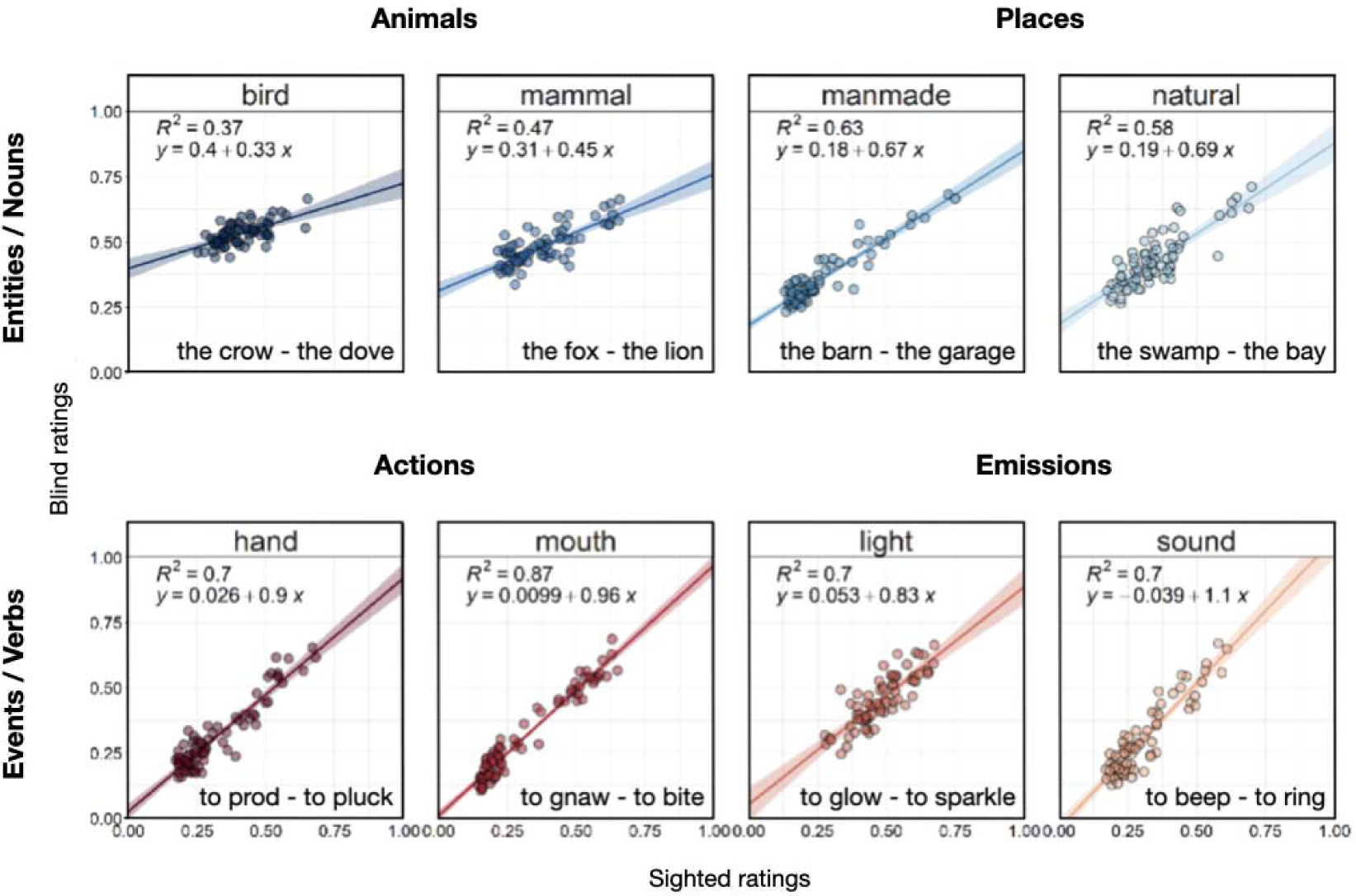
Item-wise correlations (Spearman’s rho _ρ_) between blind and sighted average group ratings. Confidence intervals (95%) are indicated via shading.

To facilitate MVPA decoding analyses, we created two non-overlapping subsets of words that were exclusively presented in either even or odd runs. This enabled us to train the classifier on one set of words and test it on a different set of words, ensuring that any above-chance classification effects reflect differences in the neural patterns associated with semantic categories and not word forms. Words in each semantic category were divided into two non-overlapping sets of 9 words. Within each set, we created all the possible pairs within a category (e.g., ‘the seagull – the parrot’, 36 pairs per set per category). There were no cross-category pairs.

### Behavioral data analysis

Due to a response box malfunction, 19/21 blind and 19/22 sighted participants contributed to behavioral data analysis. In-scanner similarity judgments were first standardized (z-scored to mean=0 ± 1SD) within each participant to account for individual differences in Likert scale use, and then standardized within grammatical class (i.e., events/verbs, entities/nouns) to a [0,1] range (i.e.,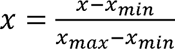). To assess agreement in semantic judgments across blind and sighted groups, we correlated item-wise ratings within each semantic category using Spearman’s rho (ρ) rank correlations. This analysis asks whether blind and sighted participants agree regarding which pairs within a semantic category are most similar in meaning (see Supplementary Materials for details).

### fMRI data acquisition

MRI structural and functional data of the whole brain were collected using a 3 Tesla Phillips scanner with a 32-channel head coil. We collected T1-weighted 3D-MPRAGE structural images using a pulse sequence in 170 sagittal slices with 1mm isotropic voxels (TE/TR=7.0/3.2ms, FoV=240×240 mm, 288×272 acquisition matrix, scan duration=5:59’). We collected T2*-functional BOLD images using parallel transverse ascending echo planar imaging (EPI) sequences in 36 axial slices with 2.5 x 2.5 x 2.5 mm voxels (TE/TR=30/2000ms, FoV=192×172mm, 76×66 acquisition matrix, 0.5mm gap, flip angle=70°, scan duration=8:04’).

### fMRI data analysis

#### Univariate analysis

Prior to preprocessing and analyzing the data, we trimmed the first 4 TRs of runs 6-8 in one congenitally blind participant (CB_24). This was necessary to remove a sharp, abrupt movement at the beginning of the runs that prevented the correct alignment of structural and functional images.

Data were analyzed using FSL, Freesurfer, the Human Connectome Project workbench, and custom in-house software written in Python (Dale, Fischl, & Sereno, 1999; Smith et al., 2004; Glasser et al., 2013). Functional data were motion corrected using FSL’s MCFLIRT algorithm (Jenkinson et al., 2002), high pass filtered to remove signal fluctuations at frequencies longer than 128 seconds/cycle, mapped to the cortical surface using Freesurfer, spatially smoothed on the cortical surface (6mm FWHM Gaussian kernel), and prewhitened to remove temporal autocorrelation. Covariates of no interest were included to account for confounds related to white matter, cerebral spinal fluid, and motion spikes.

Each of the noun and verb categories was entered as a separate predictor in a general linear model (GLM) after convolving with a canonical hemodynamic response function and its first temporal derivative. Each run was modeled separately, and runs were combined within-subject using a fixed-effects model (Dale et al., 1999; Smith et al., 2004). Group-level random-effects analyses were cluster-corrected at p<0.01 family-wise error rate (FWER) using a nonparametric permutation test.

#### ROI definition

We defined individual subjects’ functional ROIs in a two-step procedure: we first defined search spaces and then within the search spaces we defined individual-subject ROIs. Group-level search-spaces included left hemisphere regions that responded more to nouns vs. verbs or vice versa in a whole-cortex analysis (p < .05 uncorrected) and were also found to respond more to event or entity words in previous studies (see Crepaldi et al., 2013, for a review). We defined 4 noun/entity-preferring search spaces: left precuneus (LPC), left inferior parietal lobule (LIP), left lateral inferior temporal cortex (LlatIT), and left medial ventral temporal cortex (LmedVT) and 1 verb/event-preferring search-space: left middle temporal gyrus/inferior parietal cortex (LMTG+) (Supplementary Figure 5). Although the left inferior frontal gyrus also responded more to verbs than nouns, we previously found that it showed weak and category-invariant decoding in sighted adults (Elli et al., 2019); therefore, we did not use this ROI in the current study. Search spaces were first defined in the blind and sighted groups separately and then combined across groups, such that each search space (e.g., blind LPC + sighted LPC) included all the voxels responding more to verbs or nouns in either group. This procedure is inclusive to avoid omitting above-threshold activation in either of the groups.

Next, we defined individual-subject ROIs within each search space by selecting every participant’s top 300 active vertices for the verbs>nouns (verb ROI) or nouns>verbs (noun ROIs) contrasts (see Supplementary Materials for details).

Following past work demonstrating occipital activation during language processing in blind individuals (Röder et al., 2002; Amedi et al., 2003; Bedny et al., 2011, 2012, 2015; Lane et al., 2015), we additionally defined in each participant two ROIs in occipital cortex: left and right V1-V2 (BA17-18) from the PALS-B12 Brodmann area atlas included in FreeSurfer (Van Essen, 2005).

#### MVPA ROI analysis

We used MVPA (PyMVPA toolbox; Hanke et al., 2009) to assess the extent to which patterns of activity in entity- and event-responsive ROIs encode differences in semantic category within each grammatical class.

For each ROI in each participant, we trained a linear support vector machine (SVM) classifier to separately decode among the 4 verb categories and the 4 noun categories (chance 25%). We submitted to analysis the z-scored beta parameter of the GLM associated with each vertex for each semantic category in each run (2 grammatical classes * 4 categories per class = 8 total observations per run) (see Supplementary Materials for details). Within each of the entity- and event-responsive ROIs, we used one-tailed Student’s t-tests to test the classifier’s accuracy against chance (25%), and two-tailed independent samples Student’s t-test to compare the accuracy for verbs and nouns. We used repeated measures ANOVAs to test for interactions between groups, ROIs, and grammatical class (nouns/verbs). We evaluated significance using a combined permutation and bootstrapping approach (Schreiber & Krekelberg, 2013; Stelzer, Chen, & Turner, 2013) (see Supplementary Materials for details). The same approach was used to assess the statistical significance of decoding accuracies within the two occipital ROIs.

Next, to evaluate how well the classifier performed on pairwise distinctions among verbs and among nouns (e.g., birds vs. mammals), we inspected the confusion matrices generated by the classifier. The confusion matrices yield the classification and misclassification frequencies for any pair of categories, which can be compared using a signal detection theory framework (Swets, Tanner Jr, & Birdsall, 1961; Green & Swets, 1966; Haxby, Connolly, & Guntupalli, 2014). Within each ROI, we assessed the discriminability between 1) animal vs. place nouns across the entity-responsive network and 2) light verbs vs. all other verb categories in the LMTG by computing the nonparametric estimate of discriminability (Pollack & Norman, 1964; Grier, 1971; Stanislaw & Todorov, 1999). An A′ of 0.5 corresponds to chance performance, whereas 1.0 indicates perfect discriminability. Because A′ values did not follow a normal distribution, we used one-sample Wilcoxon signed rank tests to compare A′ values to chance performance and a repeated measures permutation ANOVA (5,000 permutations) using the permuco package in R (Frossard & Renaud, 2021) to test for interactions between groups, ROIs, and classification error type in entity-responsive brain regions. Wilcoxon signed rank tests use the test statistic V, which represents the sum of the positive ranks, or the distance of all observed values greater than the chance-level from the chance-level.

## Results

### Behavioral results

The similarity judgments of the blind and sighted people were significantly correlated across groups for every semantic category, but some categories were more similar across groups than others (Figure 1). Correlations across groups were highest for mouth verbs (ρ^2^=0.87), and lowest for birds (ρ^2^=0.37) and mammals (ρ^2^=0.47). When we measured coherence among individuals within a group, we likewise observed some between-group differences. An ANOVA comparing within-group agreement revealed a semantic category by group interaction, whereby blind and sighted participants’ judgments differed more within-group for some semantic categories compared to others (Supplementary Figure 2, nouns group x noun semantic category interaction, F_(3,111)_=3.47, p<0.02; group x verb semantic category interaction F_(3,111)_=2.79, p=0.04). Specifically, there was lower agreement for birds and mammals among blind than sighted people (birds: blind ρ=0.19 ± 0.18 SD; sighted ρ=0.44 ± 0.19; mammals: blind ρ=0.38 ± 0.22; sighted ρ=0.62 ± 0.14).

Birds and mammals were also the categories for which there was the largest difference in average similarity judgments between blind and sighted people. That is, people born blind tended to rate birds and mammals as more similar to each other than people who are sighted (marginal group x semantic category interaction, F_(3,111)_=2.58, p=0.06). Relative to the sighted, blind participants also provided higher noun similarity ratings overall (Supplementary Figure 1A; repeated measures ANOVA, 2 groups (sighted, blind) x 4 noun semantic categories (birds, mammals, manmade pl., natural pl.): main effect of group, F_(1,37)_=7.46, p=0.01). There were no group effects or group by condition interactions in reaction time data (Supplementary Figure 1B; see Supplementary Materials for details). These results are consistent with the notion that vision influences within-category semantic similarity judgments of animals.

### fMRI results

#### Selective responses to animals in precuneus do not require visual experience

Both groups exhibited preferential univariate responses to nouns over verbs in parietal and temporal regions previously associated with concrete entities, including the posterior parietal, lateral inferior temporal, and medial occipito-temporal cortices as well as the precuneus (Supplementary Figure 3). A whole-cortex univariate contrast between animals and places revealed that animals (birds and mammals) activated a dorsal sub-region of the PC in both sighted and blind participants (Figure 2A sighted; B blind). The dorsal animal-selective response observed in the blind group overlapped with previously reported responses to people in dorsal PC of sighted participants (e.g., Fairhall et al., 2013b). This result suggests that the emergence of a preferential response to living things in the dorsal PC does not require visual experience.

**Figure 2:**
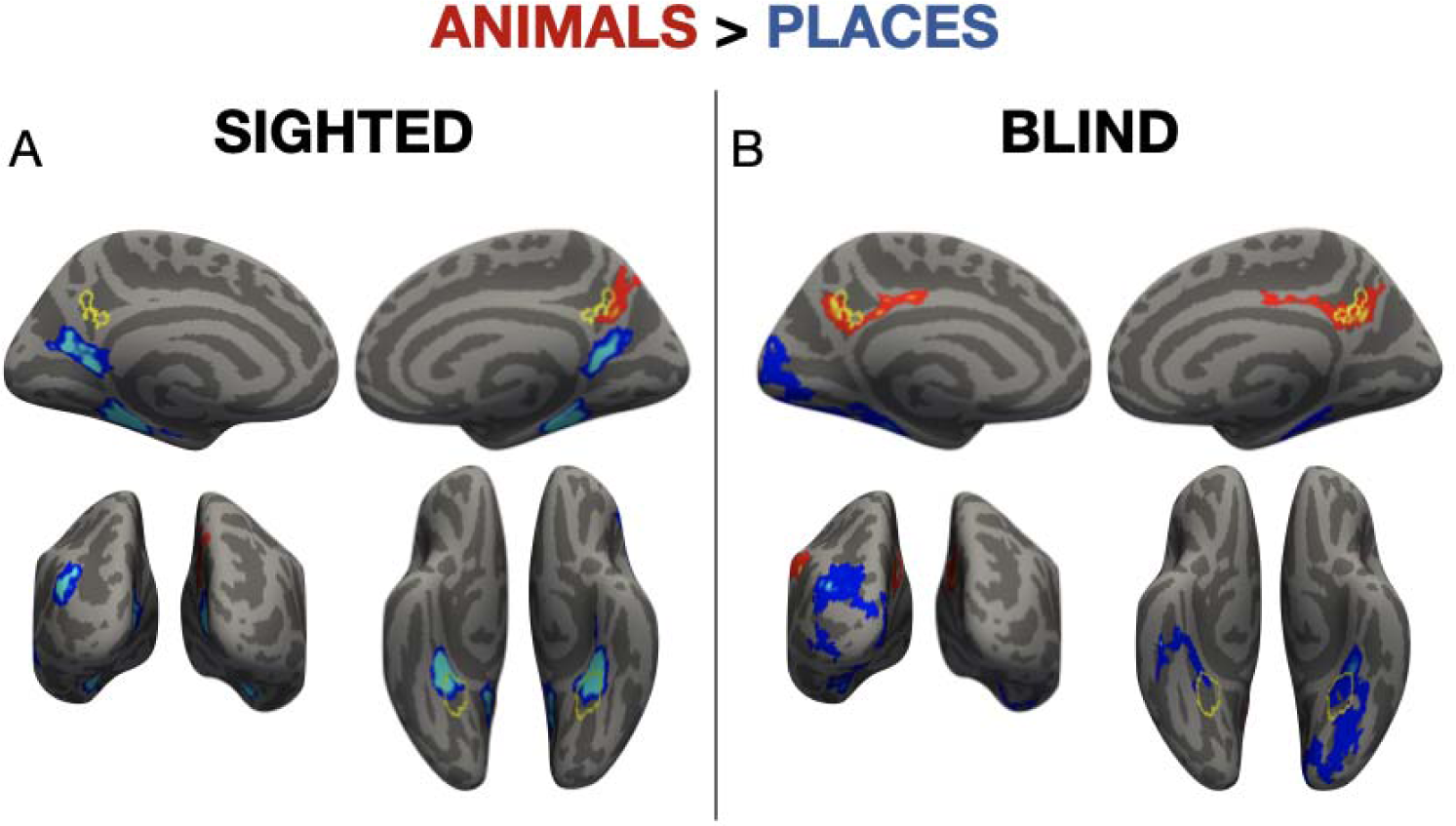
Whole-cortex results for animals>places: (A) Sighted; (B) Blind. Group maps are shown p<0.01 with FWER cluster-correction for multiple comparisons. Voxels are color coded on a scale from p=0.01 to p=0.00001. The average PPA location from separate cohort of sighted subjects (Weiner et al., 2017) is overlaid on the place noun response observed in the current study. The two overlap in both groups, with the focus of the place noun response located more anteriorly. The average people-preferring precuneus location from a separate cohort of sighted subjects (Fairhall & Caramazza, 2013b) is overlaid on the animals response observed in the current study. These also overlap in both blind and sighted participants. Increased activation for animals over places is observed in the left precuneus in sighted participants at a lower statistical threshold (p < 0.05 uncorrected). See Supplementary Figure 3 for full whole-cortex results.

Consistent with prior findings, preferential responses to place nouns were observed in an inferior portion of PC, i.e., retrosplenial complex, in sighted participants, and emerged in the same location at a more lenient statistical threshold in the blind group (p<0.01, uncorrected). This retrosplenial complex region has previously been identified as part of the ‘place’ processing network in sighted participants (Ino et al., 2002; Rauchs et al., 2008; Epstein, 2008; Dilks et al., 2022). In both groups, preferential responses to places were also observed in medial VOTC, near but somewhat anterior to the canonical location of the parahippocampal place area (PPA) (Weiner et al., 2017). PPA-like activation in the blind group extended posteriorly into early visual cortices. The same medial VOTC region showed a larger preference for places over animals in the sighted group relative to the blind group (group-by-condition interaction, Figure 5B). The only other group-by-condition interaction for the animals vs. places contrast was observed in early visual areas and is discussed in detail below.

#### Decoding of animals vs. places throughout entity responsive network is similar across blind and sighted groups

MVPA revealed that animals were discriminable from places throughout entity-responsive regions in both sighted and blind participants (all ps < .05), including in PC (sighted: V = 210, p = 0.004, blind: V = 173, p = 0.007; see Supplementary Figure 6 and Supplementary Table 4 for results in each ROI). Discriminability between animals and places was not different across ROIs, but the sighted group exhibited higher discriminability overall (repeated measures ANOVA, 2 groups (sighted, blind) x 4 ROIs (LPC, LIP, LlatIT, LmedVT): main effect of group F_(1,41)_=37.30, permuted p = 0.002; main effect of ROI F_(1,41)_=1.55, permuted p = 0.2).

Inspection of the confusion matrices showed that in both groups, birds were more likely to be confused with mammals than with places (repeated measures ANOVA, 2 groups (sighted, blind) x 2 error types (bird-mammal, bird-place) x 4 ROIs (LPC, LIP, LlatIT, LmedVT): main effect of error type F_(1,41)_=32.04, permuted p = 0.0002), although this effect was smaller in the blind than in the sighted (error type x group interaction F_(1,41)_=10.60, permuted p = 0.003). Similarly, mammals were more likely to be confused with birds than with places (repeated measures ANOVA, 2 groups (sighted, blind) x 2 error types (bird-mammal, bird-place) x 4 ROIs (LPC, LIP, LlatIT, LmedVT): main effect of error type F_(1,41)_=28.25, permuted p = 0.0002), and this effect was also smaller in the blind than in the sighted (error type x group interaction F_(1,41)_=23.92, permuted p = 0.0002). These results demonstrate the robust dissociation between animals and places in the entity-responsive network of blind people.

#### Responses to visual ‘sparkle’ verbs in LMTG+ are similar across blind and sighted people

A univariate whole-cortex analysis revealed a preference for verbs over nouns in left middle temporal gyrus (LMTG+) of blind and sighted participants (Figure 3). In a more sensitive individual-subject ROI analysis, there were no differences across groups in the LMTG+ response to light verbs or any other verb category (Figure 3; repeated measures ANOVA, 2 groups (sighted, blind) x 4 verb categories (hand, mouth, light, sound): group x verb category interaction, F_(3,123)_=1.14, p=0.34; main effect of group, F_(3,123)_=0.06, p=0.81; main effect of semantic category, F_(3,123)_=7.16, p<0.001).

**Figure 3:**
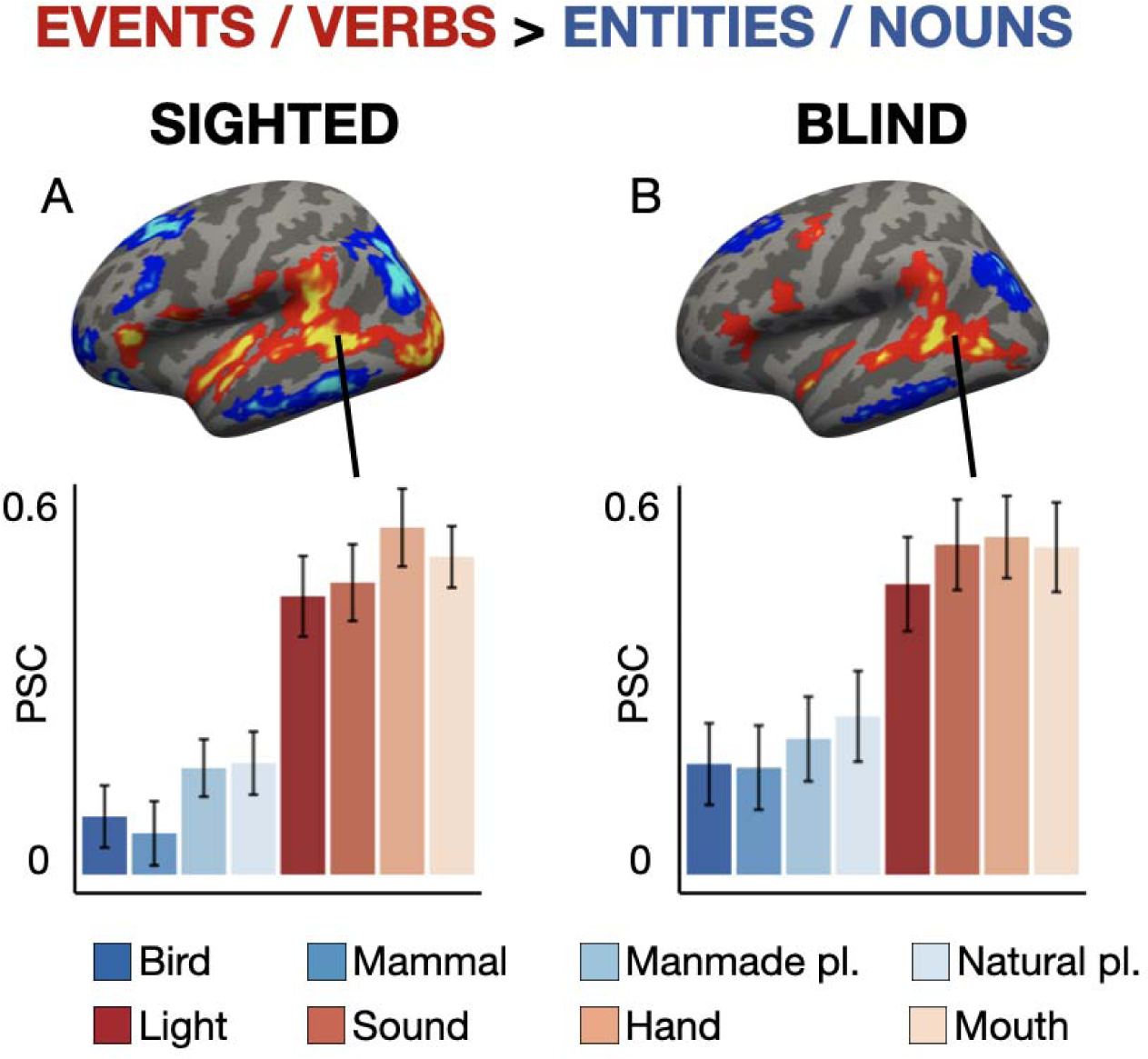
Whole-cortex results for events/verbs > entities/nouns on the left lateral surface: (A) Sighted; (B) Blind. Group maps are shown at p<0.01 with FWER cluster-correction for multiple comparisons. Voxels are color coded on a scale from p=0.01 to p=0.00001. (C) Peak percent signal change (PSC) from the 5% most active vertices for verbs>nouns in the LMTG+ (left: sighted; right: blind). Note that this figure can be used to evaluate differences among verbs and among nouns in the LMTG+ ROI, as well as differences between groups in noun/verb responses. This figure cannot be used to evaluate within group differences between verbs and nouns because the ROIs were defined as the most verb-selective vertices; thus the difference between verbs and nouns may be exaggerated due to statistical bias. See Supplementary Figure 3 for full whole-cortex results.

Consistent with the idea that the LMTG+ is sensitive to fine-grained semantic and/or grammatical distinctions among verbs, multivariate patterns of activity in LMTG+ distinguished between semantic categories of verbs in blind (t_(20)_=3.91, permuted p=0.0004) and sighted (t_(21)_=3.88, permuted p=0.0003) participants. There were no differences in decoding accuracy between the groups (repeated measures ANOVA, 2 groups (sighted, blind): main effect of group, F_(3,123)_=0.94, p=0.34; Supplementary Figure 5).

In the blind group, the LMTG+ was the only region that showed higher decoding for verbs than nouns (t_(20)_=-2.68, permuted p=0.01), providing evidence for selectivity in this population. In the sighted, decoding for verbs and nouns was not different in the LMTG+ (t_(21)_=-0.28, permuted p=0.78), whereas it was higher for nouns in LPC and LmedVT (Supplementary Figure 5; Supplementary Table 5). A 3-way repeated measures ANOVA (2 groups (sighted, blind) x 5 ROIs (LMTG+, LPC, LIP, LlatIT, LmedVT) x 2 grammatical classes (entities/nouns, events/verbs)) revealed an ROI x grammatical class interaction but no 3-way interaction with group (two-way ROI x grammatical class interaction, F_(4,164)_=6.40, permuted p<0.0001; 3-way interaction F_(4,164)_=1.31, permuted p=0.26).

Next, we compared classifier error patterns to probe the ‘representational space’ of the LMTG+ across groups. Consistent with the idea that the LMTG+ of blind and sighted people shares a similar representational space, the confusion matrices for the blind and sighted groups were significantly correlated (Figure 4B; r(30)=0.55, p=0.03). This correlation suggests the same verb categories that are more similar and thus more confusable for the sighted group are also more similar and thus more confusable for the blind group.

**Figure 4:**
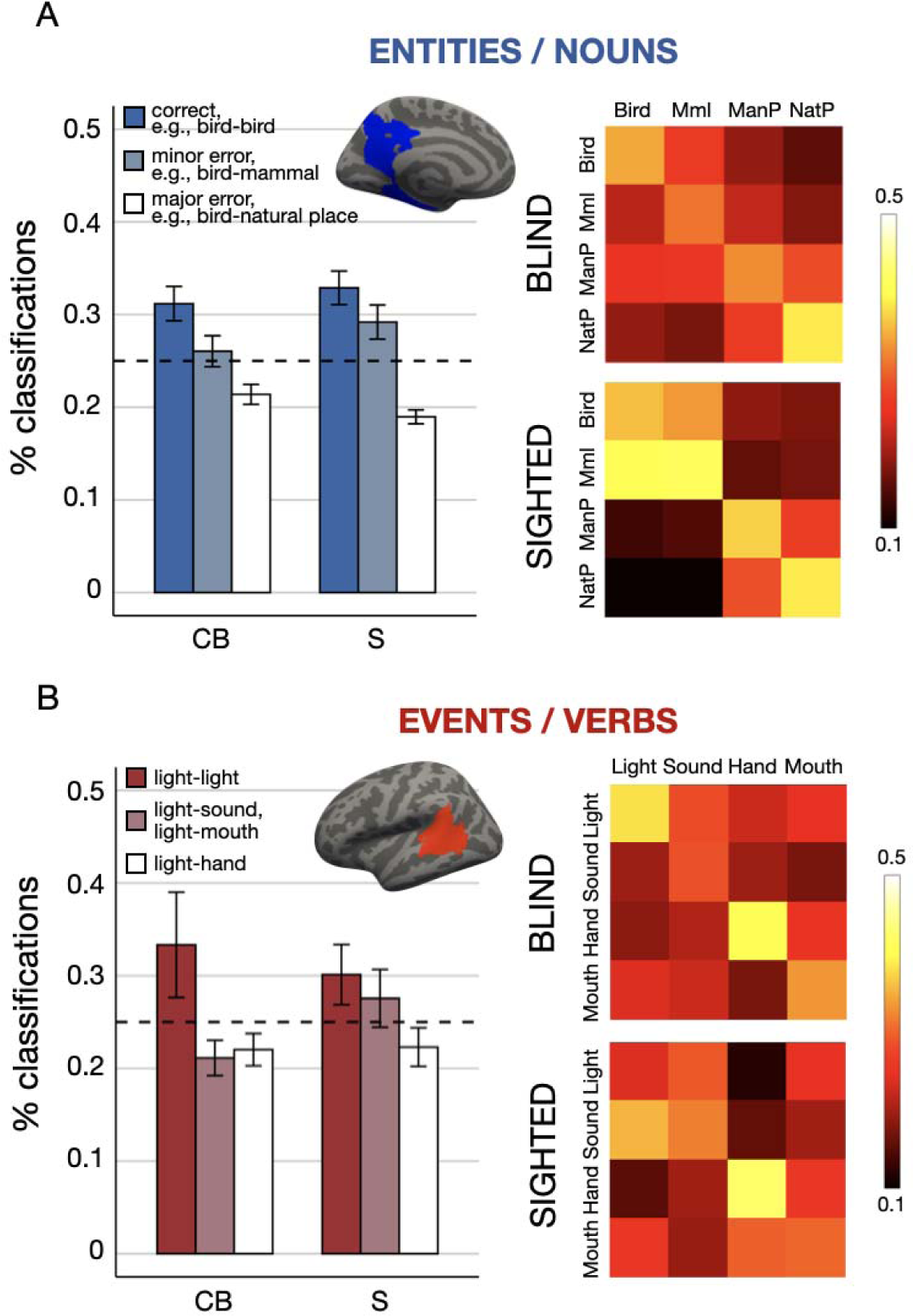
Classifier responses and confusion matrices for entity categories in the LPC (A) and for event categories i the LMTG+ (B). Bar graphs display the correct responses and errors for classification of animals vs. places (LPC) and light vs. all other verb categories (LMTG+) within each participant group. Note that in the two lightest bars reflect the number of errors made in both directions (e.g., “light-sound” = mean of light (real) – sound (predicted) and sound (real) – light (predicted)). Chance: 25%. Confusion matrices (columns = real, rows = predicted) displa the percentage of correct responses (diagonals) and errors (off diagonals) for classification of the relevant categories in each ROI. See Supplementary Figure 6 for results from all ROIs. Key: Mml = mammal, ManP = manmade place, NatP = natural place.

**Figure 5:**
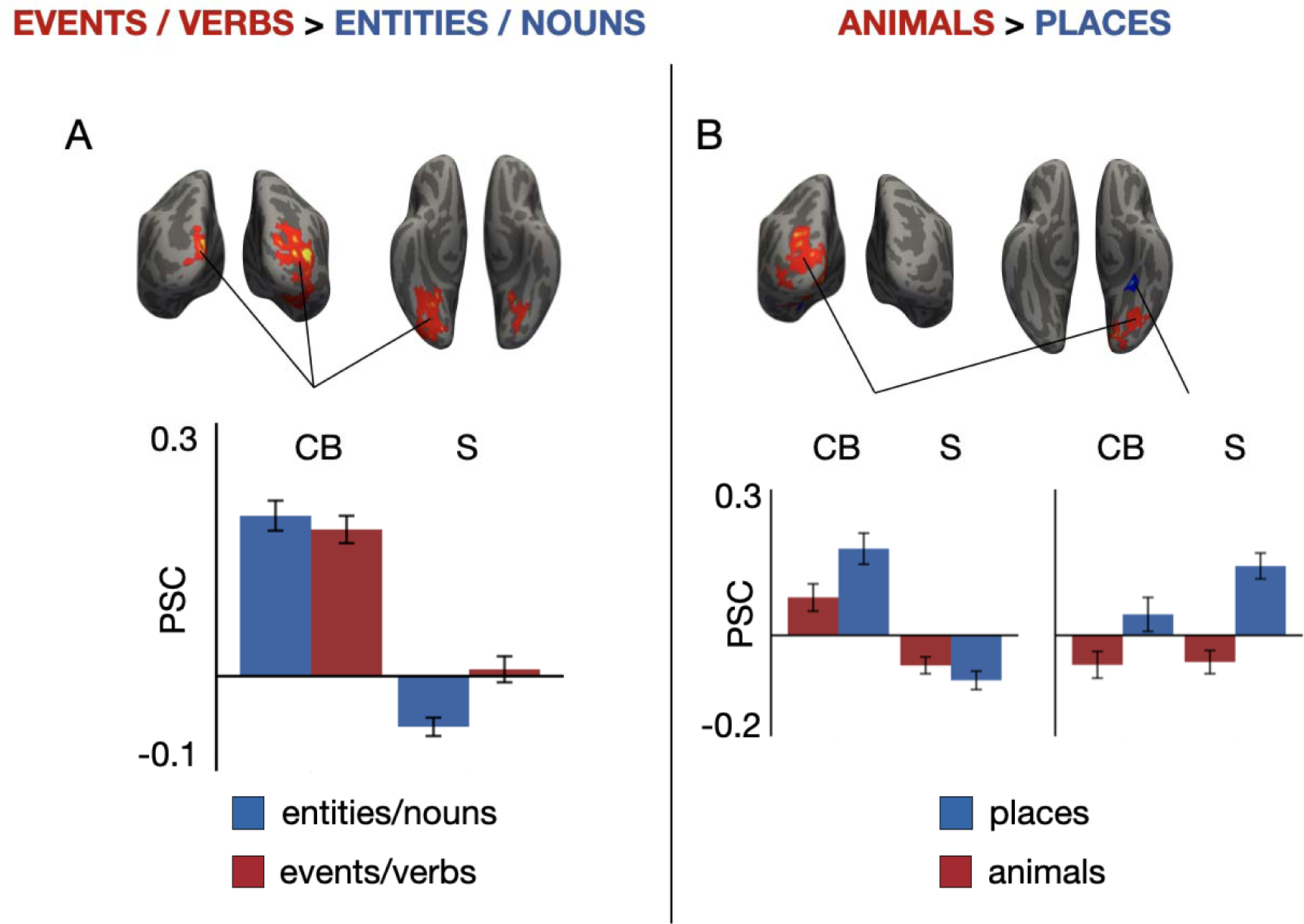
Group-by-condition interactions of univariate responses in occipital cortices. Group maps are shown p<0.01 with FWER cluster-correction for multiple comparisons. Voxels are color coded on a scale from p=0.01 t p=0.00001. (A) Peak percent signal change averaged across all occipital regions in which group-by-grammatical class (events vs. entities) interactions were observed. (B) Peak percent signal change in occipital regions in which group-by-noun category (animals vs. places) interactions were observed (occipital pole, anterior medial VTC).

Finally, we examined the classification of visual verbs separately. We found that light verbs could be distinguished from hand verbs (blind: V = 158, p = 0.0009, sighted: V = 170, p = 0.0001) in both groups. In the blind group, light verbs were also distinguishable from both sound verbs (V = 92, p = 0.007) and mouth verbs (V = 127, p = 0.009). In the sighted group, light verbs were not distinguishable from sound verbs (V = 86, p = 0.65) and were marginally distinguishable from mouth verbs (V = 113, p = 0.046). Together, these results suggest that visual verbs are distinguishable from other verb categories in the LMTG+ of both blind and sighted people.

In sum, in both sighted and blind participants, the LMTG+ responds more to verbs than nouns and distinguishes among different semantic categories of verbs, including between light verbs and other verb types.

#### Differences in responses to words in occipital networks of blind and sighted people

Group differences in univariate responses to verbs vs. nouns emerged exclusively within occipital cortices. On the medial, ventral, and dorsal surfaces of the occipital pole, particularly in the right hemisphere, sighted participants exhibited greater deactivation for nouns compared to verbs, whereas blind participants exhibited equivalent above-baseline activity for both nouns and verbs (Figure 5A). Group differences in univariate responses to animals vs. places also emerged in occipital cortices. On the medial, ventral, and dorsal surfaces of the left occipital pole, blind participants exhibited increased responses to places over animals, with above-baseline responses observed for both word types, whereas sighted participants exhibited deactivation for both animals and places (Figure 5B). As discussed above, a small region in the medial VOTC also showed a larger preference for places in the sighted. These results suggest that unlike frontal, temporal and parietal cortices, occipital cortices do not show similar word category preferences across blind and sighted groups. The same occipital areas that ‘prefer’ place words in the blind group show greater deactivation for places in the sighted, while the same areas that show deactivation for entities in the sighted show above rest responses to both entities and events in the blind. Furthermore, sighted participants showed systematic deactivation in occipital areas (e.g., for entities) but no above-rest activity for any word category. By contrast, several early occipital areas showed above-rest responses to entities and events in the blind group.

Multivariate ROI analysis revealed weak decoding among entities and among events in visual cortices in both groups (Supplementary Figure 8). In low-level visual regions defined using a Brodmann area atlas (V1-V2; BA17-18), we observed above-chance decoding in the right hemisphere of blind participants (noun categories: t_(20)_=2.51, permuted p=0.009; verb categories: t_(20)_=2.33, permuted p=0.01) and marginal decoding in the left hemisphere of sighted participants for nouns (noun categories: t_(21)_=1.56, permuted p=0.07, verb categories: t_(20)_=0.45, permuted p=0.34). Despite above-rest univariate responses to verbs and nouns in early visual cortices of blind people, these regions do not appear to robustly encode finer-grained distinctions among semantic categories in either the blind or the sighted.

## Discussion

### Preserved specialization for living things in semantic network of people born blind

Vision offers an important source of information about living things, including animals (e.g., Warrington & Shallice, 1984; Farah & McClelland, 1991). Consistent with this idea, we found that within-category semantic similarity judgments for birds and mammals (e.g., ‘crow’ vs. ‘dove’) differ between blind and sighted people, more so than judgments about places (e.g., ‘swamp’ vs. ‘bay’). For birds in particular, blind and sighted judgments were least correlated, and blind participants provided numerically higher similarity ratings compared to sighted participants. This finding is consistent with prior evidence that sighted adults living in industrialized societies rely heavily on surface-level information (e.g., visual appearance) when making within category similarity judgments about living things (animals and plants). Conversely, experts and members of cultural groups that live in closer contact with nature tend to rely more on abstract causal information such as behavioral and ecological patterns (Boster & Johnson, 1989; López et al., 1997; Proffitt, Coley, Medin, 2000; Bailenson et al., 2002; Medin & Atran, 2004). An open question is whether blind and sighted people with expertise in a living things domain (e.g., ornithology) would provide more similar ratings.

Despite these behavioral differences between groups, we observed robust neural specialization for living things in temporo-parietal networks of congenitally blind adults. Multivariate analysis revealed distinct neural patterns for animals and places across entity-responsive temporoparietal areas, including left precuneus (PC), inferior parietal lobule (LIP), lateral inferior temporal cortex (LlatIT), and medial ventral temporal cortex (LmedVT). In univariate analysis, selective responses to animals emerged in the dorsal PC of both blind and sighted participants. These results parallel findings from past studies with sighted adults implicating the PC in the representation of living things, including people-related concepts and animals (Devlin et al., 2002; Fairhall & Caramazza, 2013a; 2013b; Fairhall et al., 2014; Deen & Freiwald, 2022; Peer et al., 2015; Elli et al., 2019). One prior study also identified a preference for words describing unique people vs. human face parts in PC in blind adults (Wang et al., 2016). Together, this evidence suggests that specialization for living things in PC emerges independent of vision.

Previous studies with blind participants using animal words and sounds failed to find clear evidence of ‘living-things’ specialization in people born blind (Mahon et al., 2009; He et al., 2013; Handjaras et al., 2016; Mattioni et al., 2020; Bola et al., 2022). These studies primarily focused on VOTC, which in sighted people responds primarily to images (Levy et al., 2001; DiCarlo & Cox, 2007; Hasson et al., 2002; Srihasam et al., 2014; Arcaro & Livingstone, 2017). We similarly failed to find VOTC responses to animal words among either sighted or blind people. Conversely, several recent studies using tactile faces and human voices as stimuli have reported responses in lateral VOTC of blind people (van den Hurk et al., 2017; Ratan Murty et al., 2020; Mattioni et al., 2020; Bola et al., 2022). Such responses, however, may be driven by the distinctive communicative relevance of human faces and vocalizations. This possibility is particularly plausible given that enhanced language-related responses are observed in VOTC of people born blind (Burton et al., 2002; Lane et al., 2015; Dziegiel-Fivet et al., 2021; Tian et al., 2023). By contrast, animal words used in the current study are no more communicatively or socially relevant than words for other categories, including places and events. We therefore hypothesize that temporo-parietal responses to animal words observed in the current study reflect the retrieval of conceptual information relevant to living things. Further work is needed to understand the precise function of temporo-parietal responses to animal and people words as opposed to VOTC responses to faces and voices. However, the present results clearly demonstrate that people who are born blind develop typical responses to the animal category in temporo-parietal semantic networks.

### Preserved specialization for place nouns in medial VOTC

We observed response to place nouns in medial VOTC of both sighted and blind people (Wolbers et al., 2011; He et al., 2013; van den Hurk et al., 2017; Mattioni et al., 2020). This finding corroborates past work with blind individuals, suggesting that specialized responses to place words in medial VOTC emerges independent of vision. In the sighted group, the responses we observed to place words were anterior to but overlapping with the ‘perceptual’ PPA responses typically found for images of places (Epstein & Kanwisher, 1998; Weiner et al., 2017). These place responses are more consistent with anterior memory-related and word-related responses observed in previous studies (Fairhall et al., 2014; Häusler et al., 2022; see also Baldassano et al., 2013; Silson et al., 2016). In the blind group, the medial VOTC place response was more spatially distributed and variable across participants, extending into posterior occipital cortex. It remains unclear whether PPA-like responses in people born blind encode conceptual or perceptual information.

### Preserved responses to light events in LMTG+ of people born blind

Unlike many other conceptual categories, light events (e.g., ‘sparkle’, ‘glow’) can be perceived only through the visual modality. Despite this, we observed similar neural responses to light emission verbs among sighted and congenitally blind adults. In both populations, the LMTG+ exhibits distinctive neural responses to light verbs relative to both nouns and other verb categories (e.g., hand actions). These results build on prior evidence demonstrating preserved neural representations of motion verbs in the LMTG+ of congenitally blind individuals (Noppeney et al., 2003; Bedny et al., 2012). Here we show that neural representations of visual events in LMTG+ do not require sensory experience to emerge.

### Implications for cognitive neuroscience theories of concepts

We failed to find evidence for perceptual coding of the ‘visual’ concepts (i.e., animals or light emission events) in sighted people. Responses to all spoken words in early visual areas were either below or at rest in this group. Multivariate decoding of fine-grained semantic distinctions in early occipital cortex was also weak in both groups. In general, we did not find especially large differences between blind and sighted people for ‘visual’ concepts anywhere in the cortex. This is despite the fact that participants were making subtle within category semantic judgments (e.g., ‘flash’ vs. ‘sparkle’) and that, at least for the bird category, sighted people appeared to rely partly on appearance information when making these judgments.

Some prior studies using color words as stimuli find responses to color concepts in visual areas of sighted but not blind people (Wang et al., 2020; Bottini et al., 2020). Why such effects are observed for colors but not light events or animals is not clear. According to a dual coding perspective, sensory concepts, such as colors, are dually coded in both abstract conceptual and sensory networks, whereby only the sensory networks are influenced by blindness (Laurence & Margolis, 1999; Bi, 2021). Our results support the view that seemingly sensory concepts such as *sparkle* are represented in abstract conceptual networks and that these representations are preserved in blindness. In sighted subjects, sensory responses to ‘visual’ words are highly sensitive to context and task (e.g., they are observed when participants make detailed perceptual judgments; Hsu et al., 2011; Yee & Thompson-Schill, 2016). It is possible that more vivid imagery-based tasks (e.g., imagine a sparkling pond) could reveal differences in visual cortices of sighted and blind people.

### Responses to spoken words in congenitally blind and sighted adults

Consistent with prior evidence for functional reorganization in the occipital cortices of people born blind, early occipital areas showed above-rest activity for spoken words in this population (e.g., Röder et al., 2002; Collignon et al., 2013; see Pascual-Leone et al., 2005; Merabet & Pascual-Leone, 2010; Bedny, 2017 for reviews). By contrast, activity for spoken words was below or not different from rest in the sighted. Early occipital networks of sighted and blind people also exhibited category-specific effects: posterior occipital areas with a preference for places in blind people showed no difference between animals and places in sighted people (Supplementary Figure 7).

Despite above-baseline responses to spoken words in early visual cortices of blind people, multivariate decoding of fine-grained semantic distinctions was weak in both groups. It is possible that conceptual representations are more spatially diffuse and/or less systematic in occipital cortices of blind people, leading to weaker decoding. Another possibility is that occipital cortex contributes to sentence-level rather than word-level processing in blindness (Lane et al., 2015; Kanjlia et al., 2016; Loiotile et al., 2020). One prior study found that TMS stimulation applied to the occipital pole impairs verb generation in people born blind, inducing mostly semantic errors (Amedi et al., 2004). Whether early visual networks of blind adults are behaviorally relevant to semantic processing remains an open question (see Siuda-Krzywicka et al., 2016 for an example of a related approach with sighted individuals).

## Conclusions

To our knowledge, the current study is the first to demonstrate that neural selectivity for living things (i.e., animal) concepts develops in the absence of visual experience. We also find that light events, a purely visual semantic category, elicit similar neural signatures across blind and sighted people. We conclude that for a wide range of ‘sensory’ conceptual categories, distinctive neural signatures develop independent of sensory experience.

## Acknowledgements

We would like to thank all of the blind and sighted participants, the blind community, and the National Federation of the Blind. Without their support, this study would not be possible. We would also like to thank the F.M. Kirby Research Center for Functional Brain Imaging at the Kennedy Krieger Institute for their assistance with data collection. We thank Jeffrey Bowen for assistance with statistical analyses. This work was supported by the National Institutes of Health (R01 EY027352 to M.B.) and the Johns Hopkins University Catalyst Grant (to M.B.).

