## Supplementary Materials for "Neural specialization for ‘visual’ concepts emerges in the absence of vision"

Appendix 1: Complete list of stimuli, by category.

| **Verbs** |  |  |  |
| --- | --- | --- | --- |
| **Light** | **Sound** | **Hand** | **Mouth** |
| to blink | to ring | to stroke | to choke |
| to glow | to beep | to thump | to crunch |
| to glare | to pop | to yank | to munch |
| to flame | to squeak | to pluck | to slurp |
| to beam | to bang | to clutch | to hum |
| to blaze | to boom | to whack | to gulp |
| to shine | to swish | to jab | to gnaw |
| to flash | to chime | to swat | to purse |
| to gleam | to creak | to clench | to swig |
| to flare | to shriek | to prod | to consume |
| to glint | to clink | to pelt | to whistle |
| to sparkle | to plop | to flog | to mutter |
| to glitter | to crackle | to caress | to nibble |
| to twinkle | to rattle | to pummel | to guzzle |
| to flicker | to sizzle | to fondle | to gobble |
| to glimmer | to clatter | to wallop | to pucker |
| to shimmer | to jingle | to clobber | to bite |
| to glisten | to fizzle | to snag | to lick |
| **Nouns** |  |  |  |
| **Birds** | **Mammals** | **Manmade places** | **Natural places** |
| the crow | the deer | the barn | the creek |
| the duck | the goat | the shed | the pit |
| the dove | the fox | the maze | the gulf |
| the swan | the calf | the dam | the swamp |
| the goose | the ram | the lounge | the reef |
| the fowl | the moose | the booth | the ridge |
| the crane | the bull | the vault | the marsh |
| the hawk | the ox | the mosque | the brook |
| the finch | the elk | the rink | the dune |
| the quail | the sheep | the shrine | the canyon |
| the owl | the sloth | the shack | the iceberg |
| the falcon | the boar | the moat | the glacier |
| the parrot | the hog | the castle | the meadow |
| the seagull | the giraffe | the dungeon | the crater |
| the peacock | the lion | the igloo | the boulder |
| the sparrow | the hippo | the tepee | the prairie |
| the vulture | the rhino | the garage | the bay |

Appendix 2: Matching of word stimuli

Words were matched across semantic categories in syllable length (based on the CMU Pronouncing Dictionary; Weide, 1998), phonological neighborhood size (using N-Watch; Davis, 2005), familiarity, and concreteness based on ratings collected on Amazon Mechanical Turk (AMT) (see for details Elli et al., 2019). We did not match nouns and verbs on imageability. Verbs are inherently less imageable than nouns (Bird et al., 2001; 2003). Matching them would therefore result in unrepresentative class members and make it impossible to study the most ‘visual’ nouns (living things) and verbs (light emission), which was the goal of the current study. A separate group of AMT participants also rated all possible pairs of words in the experiment according to their semantic similarity to verify that entities/nouns and events/verbs did not differ from each other in their overall semantic distance (Elli et al., 2019).

Appendix 3: Analysis of semantic similarity judgments

To compare group agreement across semantic categories, we averaged ratings for a given pair across participants within a group and then correlated ratings across groups. To measure the variability among participants within groups, we computed within-group coherence (i.e., the agreement among blind and among sighted participants) by correlating each participant to their group mean in a leave-one-participant-out correlation procedure (i.e., $\rho=\text{corr}\left( \text{s}_{\text{i}}\text{, N}-\text{s}_{\text{i}} \right)$). We tested whether blind and sighted groups differed across semantic categories in within-group coherence using two-tailed independent samples Student’s t-tests and ANOVAs on Fisher-Z transformed single-subjects’ ρ values. All Spearman’s rank correlations were computed on participants’ standardized ratings using the Hmisc package in R (Harrell & Dupont, 2014).

Appendix 4: Multi-voxel pattern analysis

For each ROI in each participant, we trained a linear support vector machine (SVM) classifier to separately decode among the 4 verb categories and the 4 noun categories (chance 25%). We obtained the z-scored beta parameter of the GLM associated with each vertex for each semantic category in each run (2 grammatical classes * 4 categories per class = 8 total observations per run). To eliminate run effects, we then normalized (mean=0, SD=1) the z-scored beta values assigned to each vertex with respect to the mean signal for that vertex across all 8 semantic categories. This normalization procedure was carried out separately within each run, before restricting the dataset to either the verb or the noun categories, depending on the analysis (i.e., MVPA for verb and noun categories was conducted separately).

We used a two-folds (even/odd) cross-validation split: the classifier was trained on half of the data (e.g., even runs) and tested on the other half (e.g., odd runs). Classification accuracy was then averaged across the two even/odd splits. Note that since even and odd runs contained different verb and noun pairs, the classifier was trained and tested on different, non-overlapping subsets of words.

Within each of the entity- and event-responsive ROIs, we used one-tailed Student’s t-tests to test the classifier’s accuracy against chance (25%), and two-tailed independent samples Student’s t-test to compare the accuracy for verbs and nouns. We used repeated measures ANOVAs to test for interactions between groups, ROIs, and grammatical class (nouns/verbs). We evaluated significance using a combined permutation and bootstrapping approach (Schreiber & Krekelberg, 2013; Stelzer, Chen, & Turner, 2013). In this approach, t- and F-statistics obtained for the observed data are compared against an empirically generated null distribution of statistical values for each test (see Elli et al., 2019 for details on the permutation testing and bootstrapping steps). We report the t- and F-values obtained for the observed data and the nonparametric permuted P-values, which correspond to the proportion of shuffled analyses that generated comparable or higher t/F values. The same approach was used to assess the statistical significance of decoding accuracies within the two occipital ROIs.

Next, to evaluate how well the classifier performed on pairwise distinctions among verbs and among nouns (e.g., birds vs. mammals), we inspected the confusion matrices generated by the classifier. The confusion matrices yield the classification and misclassification frequencies for any pair of categories, which can be compared using a signal detection theory framework (Swets, Tanner Jr, & Birdsall, 1961; Green & Swets, 1966; Haxby, Connolly, & Guntupalli, 2014). Within each ROI, we assessed the discriminability between 1) animal vs. place nouns across the entity-responsive network and 2) light verbs vs. all other verb categories in the LMTG by computing the nonparametric estimate of discriminability (Pollack & Norman, 1964; Grier, 1971; Stanislaw & Todorov, 1999). An A′ of 0.5 corresponds to chance performance, whereas 1.0 indicates perfect discriminability. Because A′ values did not follow a normal distribution, we used one-sample Wilcoxon signed rank tests to compare A′ values to chance performance and a repeated measures permutation ANOVA (5,000 permutations) using the permuco package in R (Frossard & Renaud, 2021) to test for interactions between groups, ROIs, and classification error type in entity-responsive brain regions. Wilcoxon signed rank tests use the test statistic V, which represents the sum of the positive ranks, or the distance of all observed values greater than the chance-level from the chance-level.

Appendix 5: Full results of behavioral data analysis.

*Mean ratings - nouns*

Relative to the sighted, blind participants provided higher similarity ratings overall (repeated measures ANOVA, 2 groups (sighted, blind) x 4 noun semantic categories (birds, mammals, manmade places, natural places): main effect of group, F_(1,37)_=7.46, p=0.01). Across both groups, manmade items were judged to be less similar amongst themselves compared to any other noun category (main effect of semantic category, F_(3,111)_=54.68, p<0.0001). In addition, there was a marginal interaction between group and semantic category, whereby blind participants tended to rate birds and mammals as more similar to each other compared to sighted participants (group x semantic category interaction, F_(3,111)_=2.58, p=0.06).

*Mean ratings - verbs*

Blind and sighted participants did not differ in how they rated the similarity of verbs. Blind and sighted groups judged the light emission verbs to be more similar amongst themselves than any other verb category (repeated measures ANOVA, 2 groups (sighted, blind) x 4 verb categories (hand, mouth, light, sound): main effect of semantic category, F_(3,111)_=90.42, p<0.0001; main effect of group, F_(1,37)_=2.47, p=0.13; group x semantic category interaction, F_(3,111)_=0.39, p=0.76).

*Reaction time - nouns*

Across both groups, participants were faster to make judgments about birds and mammals than manmade and natural places (repeated measures ANOVA, 2 groups (sighted, blind) x 4 noun categories (birds, mammals, manmade places, natural places): main effect of semantic category, F_(3,111)_=7.81, p<0.001). There were no group effects or group by condition interactions in RT (main effect of group, F_(1,37)_=0.53, p=0.47; group x semantic category interaction, F_(3,111)_=0.72, p=0.54).

*Reaction time - verbs*

Across both groups, participants were faster to make judgments about mouth actions than all other verb categories (repeated measures ANOVA, 2 groups (sighted, blind) x 4 noun categories (hand, mouth, light, sound): main effect of semantic category F_(3,111)_=7.37, p<0.001). There were no group effects or group by condition interactions in RT (main effect of group, F_(1,37)_=0.03, p=0.85; group x semantic category interaction, F_(3,111)_=0.67, p=0.56).

Appendix 6: Detailed explanation of the creation of functional ROIs in individual participants.

We defined subjects’ functional ROIs in a two-step procedure: we first defined search spaces and then defined individual-subject ROIs within these search spaces. First, we defined group-level search spaces in blind and sighted groups separately based on each group’s whole-cortex results (p < .05 uncorrected) for the verbs vs. nouns contrast in the left hemisphere: 4 noun/entity-preferring: left precuneus (LPC), left inferior parietal lobule (LIP), left lateral inferior temporal cortex (LlatIT), and left medial ventral temporal cortex (LmedVT) and 1 verb/event-preferring: left middle temporal gyrus, which extended in the superior temporal gyrus and inferior aspect of the parietal cortex (LMTG+). To identify LMTG, we selected a contiguous cluster of activated voxels (verb>noun) within an LMTG parcel from Fedorenko et al. (2010). To identify LPC, we selected a contiguous cluster of activated voxels (nouns>verbs) within a combination of Brodmann areas spanning the precuneus: Brodmann areas 6, 24, 35, 36, 37, 39 and 48. To identify LmedVT, we selected a contiguous cluster of activated voxels (nouns>verbs) within a combination of Brodmann areas spanning ventral occipito-temporal cortex: Brodmann areas 5, 14, 29, 30, 49. Finally, to identify LIP and LlatIT, we selected contiguous clusters of activated voxels (nouns>verbs) in the approximate location of the LIP and LlatIT search spaces used in Elli et al. (2019). To avoid bias across groups, we took the union of each group-level search space (e.g., blind LPC + sighted LPC) to create group-unbiased search spaces encompassing the activation observed in both group maps. This way, the search spaces included all voxels active above threshold in either group.

Next, we defined individual-subject ROIs within each of the group-unbiased search spaces by selecting each participant’s top 300 active vertices for the verbs>nouns (verb ROI) or nouns>verbs (noun ROIs) contrasts. These subject-specific voxels did not need to be contiguous inside the search space.

Supplementary Table 1: Participants’ demographic information

| **Participant** | **Gender** | **Age** | **Cause of blindness** | **Light** | **Education level** |
| --- | --- | --- | --- | --- | --- |
| CB_01 | F | 67 | Retinopathy of prematurity | NLP | BA |
| CB_02 | M | 47 | Unknown. Possibly an infection. | LP | BA |
| CB_04 | M | 32 | Born without optic nerve | NLP | BA - |
| CB_05 | F | 64 | Retinopathy of prematurity | NLP | HS |
| CB_06 | F | 18 | Lebers Congenital Amaurosis | LP | BA in progress |
| CB_08 | F | 29 | Retinopathy of prematurity/RLF | LP | MA |
| CB_09 | F | 65 | Retinitis Pigmentosa | LP | BA |
| CB_10 | F | 35 | Retinopathy of prematurity, Glaucoma | LP | BA |
| CB_13 | F | 52 | Lebers Congenital Amaurosis | LP | MA |
| CB_14 | M | 51 | Lebers Congenital Amaurosis | NLP | JD |
| CB_15 | F | 43 | Retinopathy of prematurity | NLP | PhD in progress |
| CB_16 | F | 31 | Optic Nerve Detached | NLP | BA in progress |
| CB_17 | F | 28 | Lebers Congenital Amaurosis | LP | BA |
| CB_18 | M | 28 | Lebers Congenital Amaurosis | LP | BA in progress |
| CB_19 | F | 34 | Weak optic nerve | LP | MA |
| CB_20 | M | 32 | Lebers Congenital Amaurosis | LP | MBA in progress |
| CB_21 | F | 25 | Retinopathy of prematurity | LP | PhD in progress |
| CB_22 | M | 32 | Anopthamalia | NLP | BA |
| CB_23 | M | 29 | Retinopathy of prematurity | NLP | Some college |
| CB_24 | F | 39 | Lebers Congenital Amaurosis |  | MA |
| CB_25 | M | 41 | Retinopathy of prematurity | NLP | BA |
| **Average** |  |  |  |  |  |
| Blind (N=21) | 13F | 39.14 | - | - |  |
| Sighted (N=22) | 16F | 37.55 | - | - |  |

Supplementary Table 2: Example stimuli

| **Entities / Nouns** | **Animals** | **Birds** | the crow – the dove | the goose – the owl |
| --- | --- | --- | --- | --- |
|  |  | **Mammals** | the fox – the lion | the giraffe – the hippo |
|  | **Places** | **Manmade** | the barn – the garage | the shrine – the temple |
|  |  | **Natural** | the swamp – the bay | the canyon – the crater |
| **Events / Verbs** | **Actions** | **Hand** | to prod – to pluck | to stroke – to pummel |
|  |  | **Mouth** | to gnaw – to bite | to slurp – to lick |
|  | **Emissions** | **Light** | to glow – to sparkle | to shine – to flash |
|  |  | **Sound** | to beep – to ring | to squeak – to bang |

Supplementary Table 3: Classifier performance against chance (25%) for verb and noun categories in LMTG+, LPC, LIP, LlatIT, and LmedVT. Permuted and Bonferroni-corrected (across ROIs) p-values are reported.

| **ROI** | **Classification accuracy** | **t** | **Permuted p** | **Bonferroni adj. p** |
| --- | --- | --- | --- | --- |
| **Sighted (n=22)** |  |  |  |  |
| **Verbs** |  |  |  |  |
| LMTG+ | 32.1% | 3.88 | 0.0003 | 0.018 |
| LPC | 27.6% | 1.14 | 0.1203 | 1 |
| LIP | 34.1% | 3.32 | 0.0007 | 0.032 |
| LlatIT | 36.1% | 4.90 | 0 | 0 |
| LmedVT | 27.7% | 1.57 | 0.0527 | 1 |
| **Nouns** |  |  |  |  |
| LMTG+ | 31.4% | 3.59 | 0.0009 | 0.018 |
| LPC | 38.2% | 4.99 | 0.0001 | 0 |
| LIP | 40.8% | 4.87 | 0.0001 | 0 |
| LlatIT | 39.1% | 3.97 | 0.0001 | 0.008 |
| LmedVT | 41.6% | 6.02 | 0 | 0 |
| **Blind (n=21)** |  |  |  |  |
| **Verbs** |  |  |  |  |
| LMTG+ | 35.1% | 3.91 | 0.0004 | 0.008 |
| LPC | 28.1% | 1.47 | 0.0726 | 1 |
| LIP | 28.9% | 1.99 | 0.0254 | 0.604 |
| LlatIT | 30.7% | 2.71 | 0.0043 | 0.134 |
| LmedVT | 30.4% | 2.29 | 0.0121 | 0.328 |
| **Nouns** |  |  |  |  |
| LMTG+ | 26.2% | 0.64 | 0.2623 | 1 |
| LPC | 34.2% | 3.29 | 0.0006 | 0.036 |
| LIP | 36.6% | 4.54 | 0 | 0.002 |
| LlatIT | 35.3% | 3.12 | 0.0023 | 0.054 |
| LmedVT | 34.7% | 3.85 | 0.0004 | 0.01 |

Supplementary Table 4: Discriminability between animals and places in LPC, LIP, LlatIT, and LmedVT, assessed using one-sample Wilcoxon signed rank tests against chance. V is the sum of the positive ranks. Uncorrected and Bonferroni-corrected (across ROIs) p-values are reported. Across the 4 noun ROIs, no differences in discriminability between animals and places were observed (repeated measures ANOVA, 2 groups (sighted, blind) x 4 ROIs (LPC, LIP, LlatIT, LmedVT): main effect of ROI F_(1,41)_=1.55, permuted p = 0.2), although the sighted group exhibited higher discriminability overall (main effect of group F_(1,41)_=37.30, permuted p = 0.002).

| **ROI** | **V** | **Uncorr. p** | **Bonferroni adj. p** |
| --- | --- | --- | --- |
| **Sighted (n=22)** |  |  |  |
| LPC | 210 | 0 | 0.0004 |
| LIP | 252 | 0 | 0.0002 |
| LlatIT | 248 | 0 | 0.0003 |
| LmedVT | 253 | 0 | 0.0002 |
| **Blind (n=21)** |  |  |  |
| LPC | 173 | 0.0009 | 0.0072 |
| LIP | 197 | 0.0003 | 0.0025 |
| LlatIT | 184 | 0.0017 | 0.0135 |
| LmedVT | 169.5 | 0.0014 | 0.0115 |

Supplementary Table 5: Comparison of classifier performance for verb vs. noun categories in LMTG, LPC, LIP, LlatIT, and LmedVT. Permuted and Bonferroni-corrected (across ROIs) p-values are reported.

| **ROI** | **Mean difference** | **t** | **Permuted p** | **Bonferroni adj. p** |
| --- | --- | --- | --- | --- |
| **Sighted (n=22)** |  |  |  |  |
| **Nouns vs. Verbs** |  |  |  |  |
| LMTG+ | -0.7% | -0.28 | 0.7812 | 1 |
| LPC | 10.7% | 3.24 | 0.0034 | 0.04 |
| LIP | 6.7% | 1.89 | 0.0685 | 0.721 |
| LlatIT | 3% | 0.74 | 0.4639 | 1 |
| LmedVT | 13.9% | 3.53 | 0.0018 | 0.02 |
| **Blind (n=21)** |  |  |  |  |
| **Nouns vs. Verbs** |  |  |  |  |
| LMTG+ | -8.9% | -2.68 | 0.0147 | 0.145 |
| LPC | 6.1% | 1.84 | 0.0827 | 0.806 |
| LIP | 7.8% | 2.5 | 0.0209 | 0.211 |
| LlatIT | 4.6% | 1.32 | 0.2007 | 1 |
| LmedVT | 4.3% | 1.13 | 0.2675 | 1 |

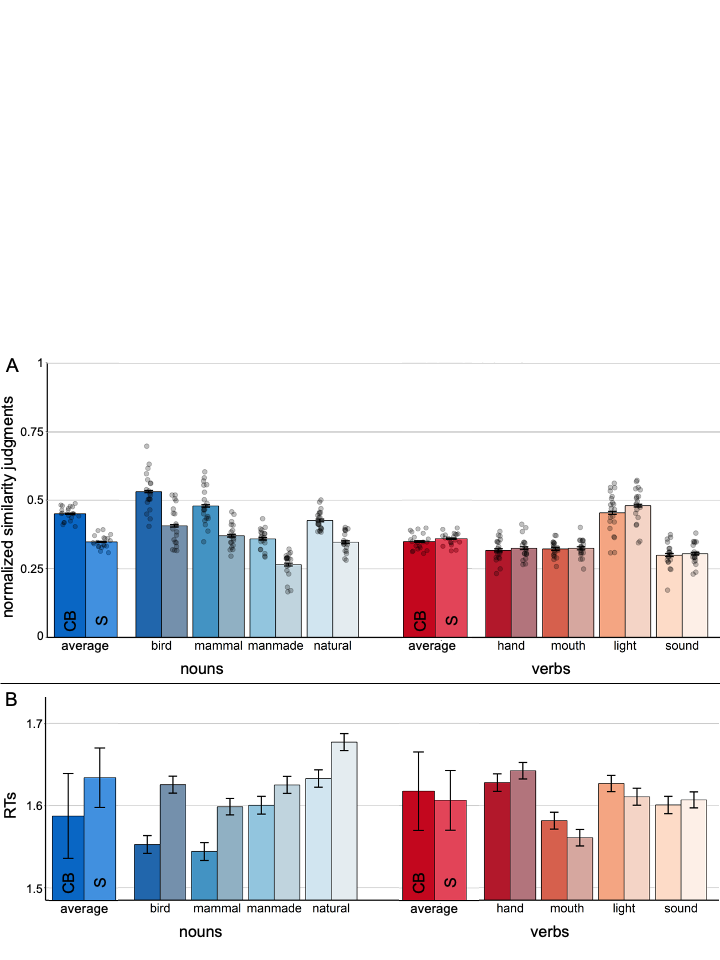

Supplementary Figure 1: In-scanner behavioral results by semantic category

: (A) Average similarity judgments; (B) reaction times. Error bars: ± standard error of the mean.

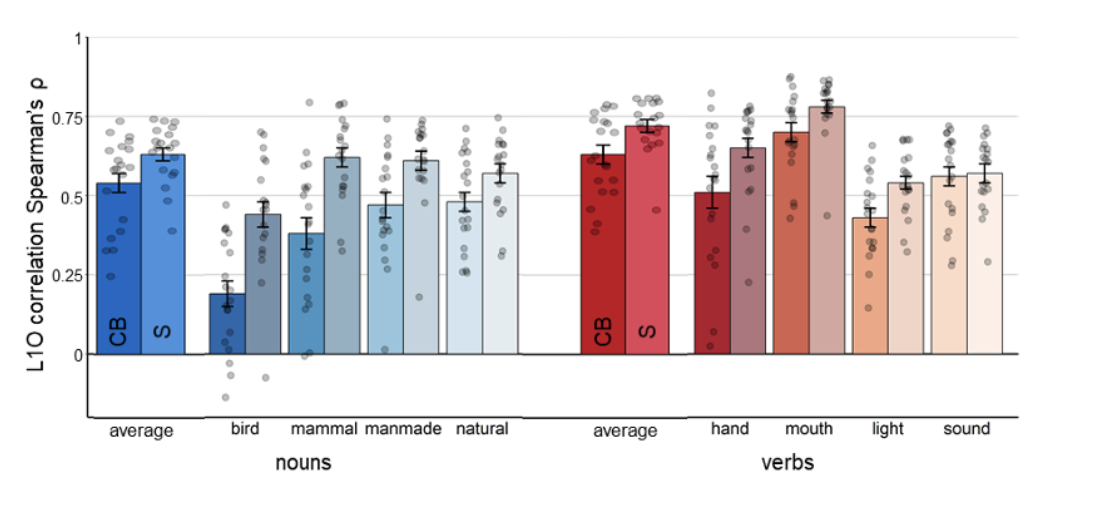

Supplementary Figure 2: In-scanner behavioral results by semantic category: Leave-one-out within-group correlations (Spearman’s ρ). Error bars: ± standard error of the mean.

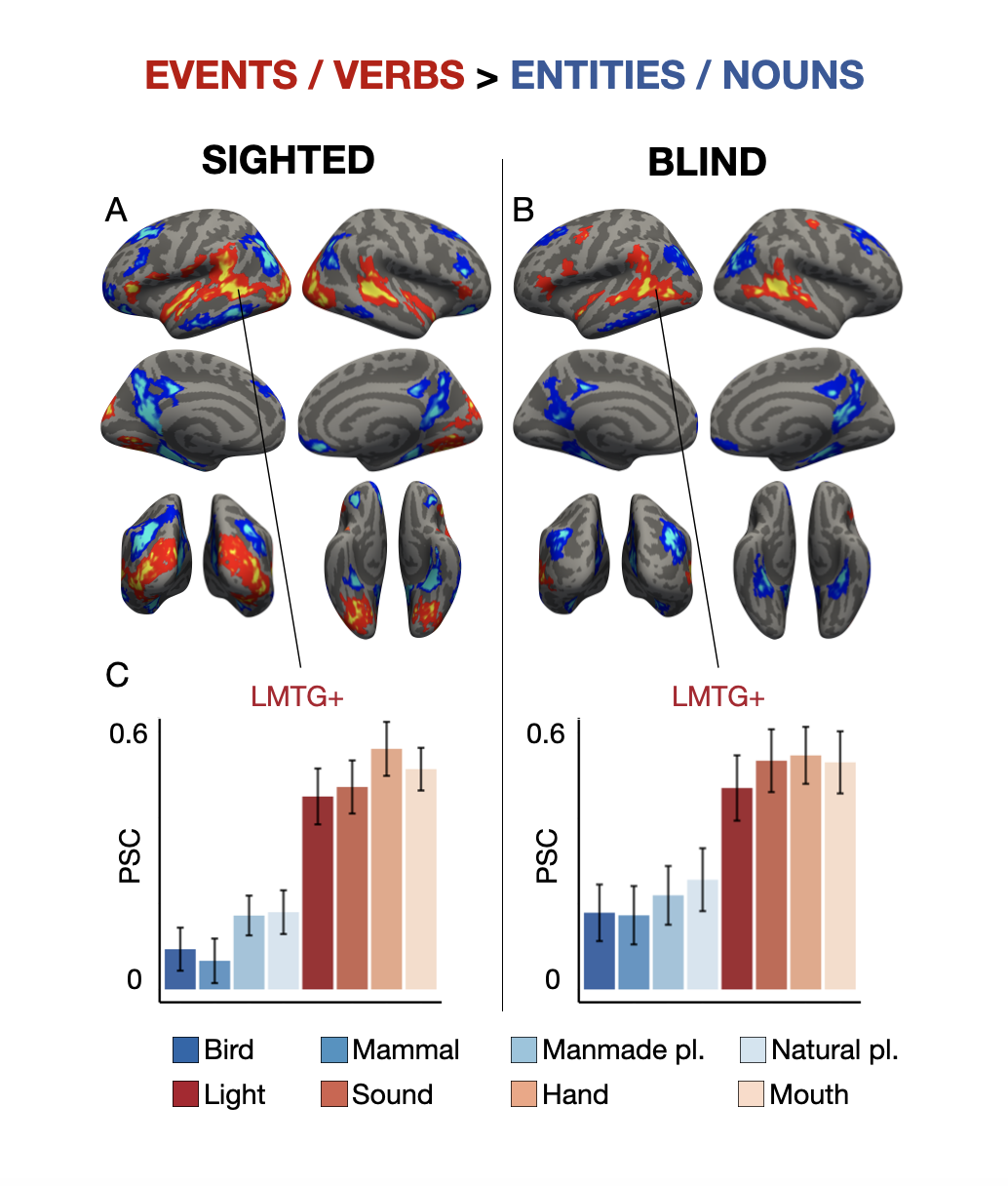

Supplementary Figure 3: Whole-cortex results for events/verbs > entities/nouns

: (A) Sighted; (B) Blind. Group maps are shown at p<0.01 with FWER cluster-correction for multiple comparisons. Voxels are color coded on a scale from p=0.01 to p=0.00001. (C) Peak percent signal change (PSC) from the 5% most active vertices for verbs>nouns in the LMTG, extending into superior temporal gyrus and inferior parietal cortex (LMTG+). Note that this figure can be used to evaluate differences among verbs and among nouns in the LMTG+ ROI, as well as differences between groups in noun/verb responses. This figure cannot be used to evaluate within group differences between verbs and nouns because the ROIs were defined as the most verb-selective vertices; thus the difference between verbs and nouns may be exaggerated due to statistical bias.

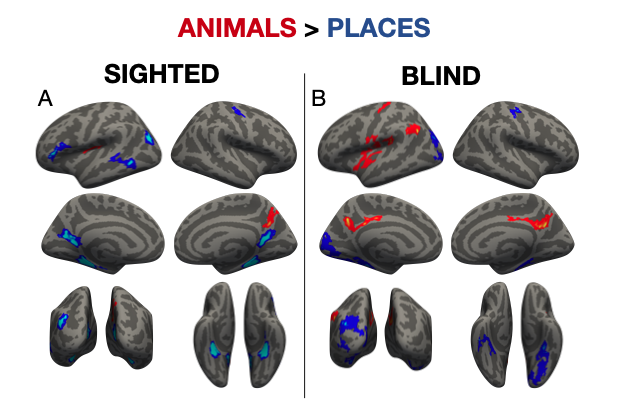

Supplementary Figure 4: Whole-cortex results for animals > places

: (A) Sighted; (B) Blind. Group maps are shown at p<0.01 with FWER cluster-correction for multiple comparisons. Voxels are color coded on a scale from p=0.01 to p=0.00001.

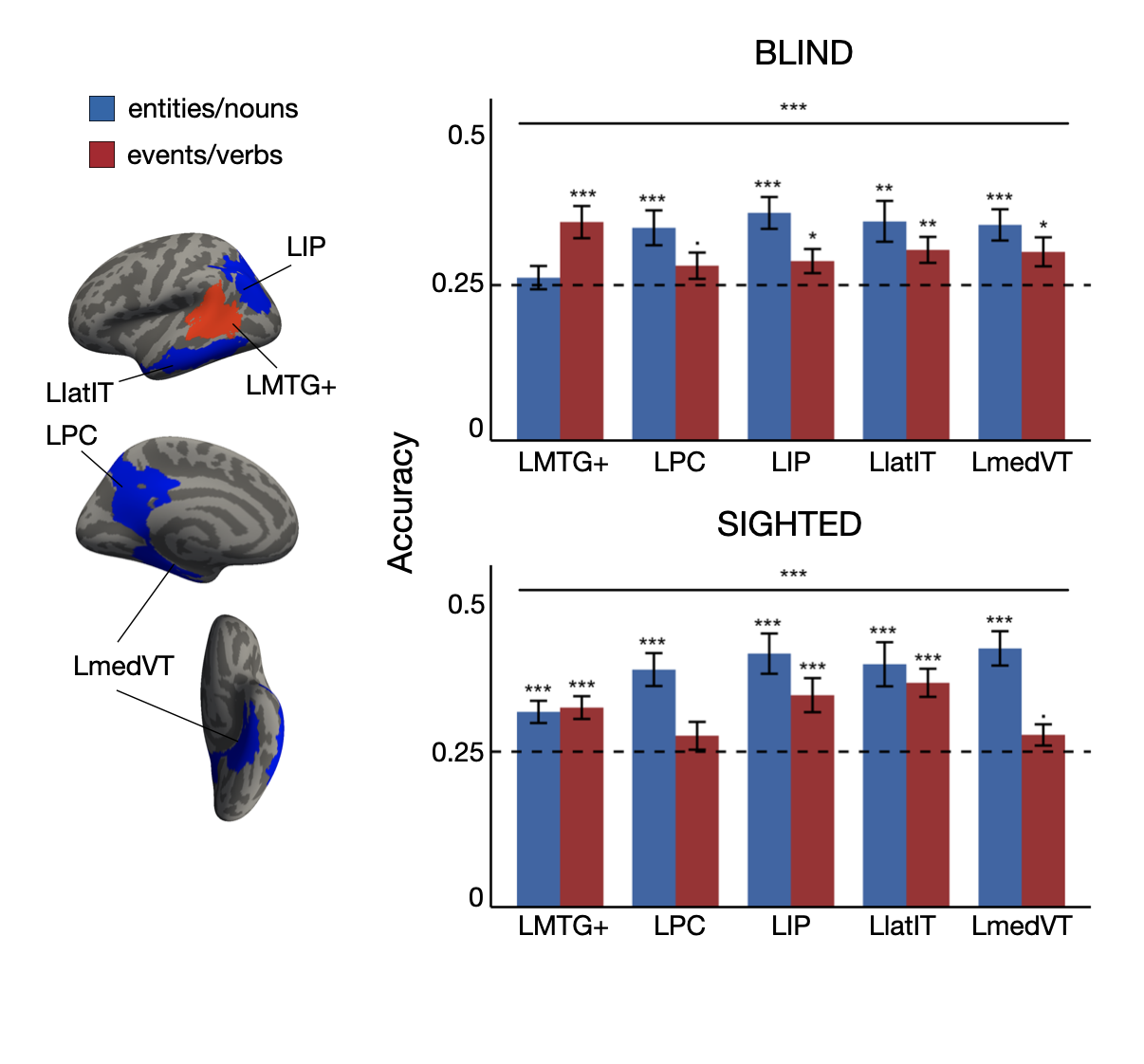

Supplementary Figure 5: Classification accuracy in verb (LMTG+) and noun (LIP, LPC, LlatIT and LmedVT) selective regions in each participant group

. Left: Group search spaces in which individual regions of interest were defined. Right: Bars representing classifier performance for each group and grammatical class. Chance: 25%. Signif. codes: 0 ‘***’ 0.001 ‘**’ 0.01 ‘*’ 0.05 ‘.’ 0.1 ‘’ 1.

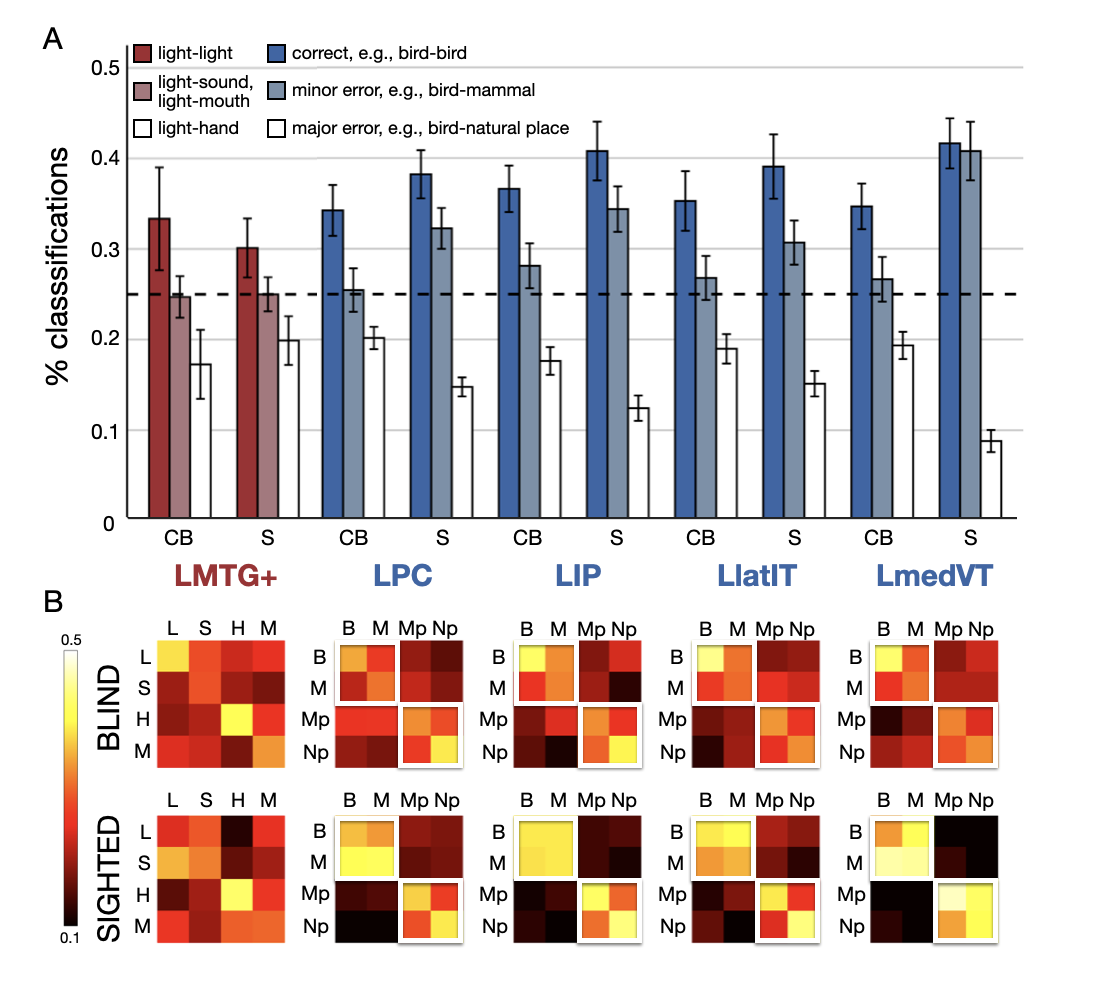

Supplementary Figure 6: Classifier responses and confusion matrices

. (A) Bar graphs display the correct responses and errors for classification of light vs. all other verb categories (LMTG+) and animals vs. places (noun ROIs) for each group and ROI. Note that the two lightest bars reflect the number of errors made in both directions (e.g., “light-sound” = mean of light (real) – sound (predicted) and sound (real) – light (predicted)). Chance: 25%. (B) Confusion matrices (columns = real, rows = predicted) display the percentage of correct responses (diagonals, highlighted in white in noun ROIs for animals vs. places) and errors (off diagonals) for classification of verb categories in LMTG+ and noun categories in LPC, LIP, LlatIT, and LmedVT. Key: L = light, S = sound, H = hand, M = mouth; B = bird, M = mammal, Mp = manmade place, Np = natural place.

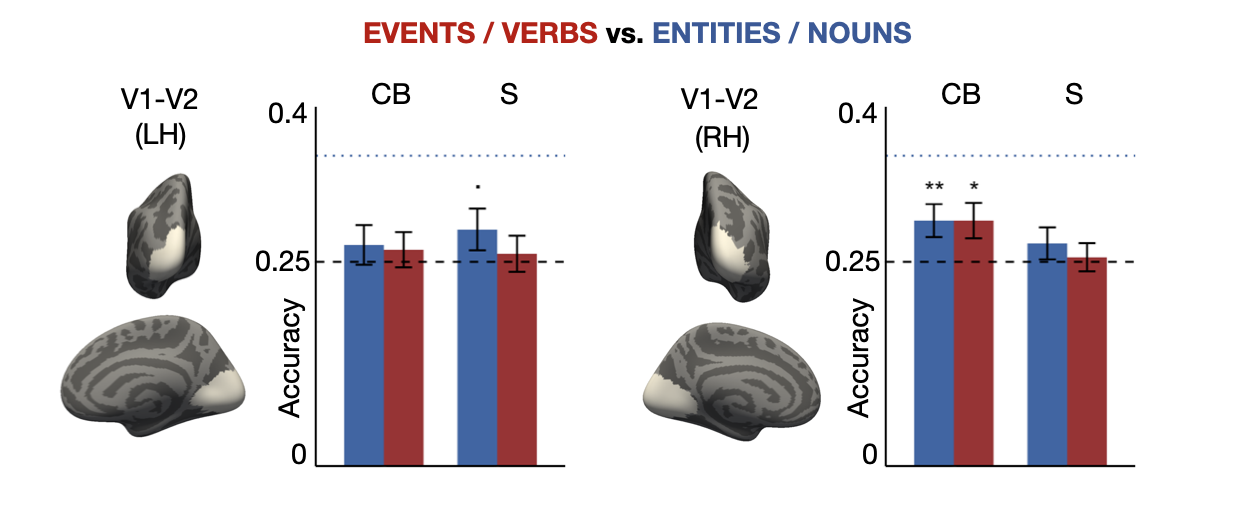

Supplementary Figure 7: Classification accuracy in each participant group for noun and verb categories in bilateral V1-V2 (BA 17-18). Chance: 25%. The blue dotted line shows the average decoding accuracy (=38%) for verb categories in the verb ROI (LMTG+) and noun categories in the noun ROIs (LIP, LPC, LlatIT and LmedVT) in the sighted group. Signif. codes: 0 ‘***’ 0.001 ‘**’ 0.01 ‘*’ 0.05 ‘.’ 0.1 ‘’ 1.

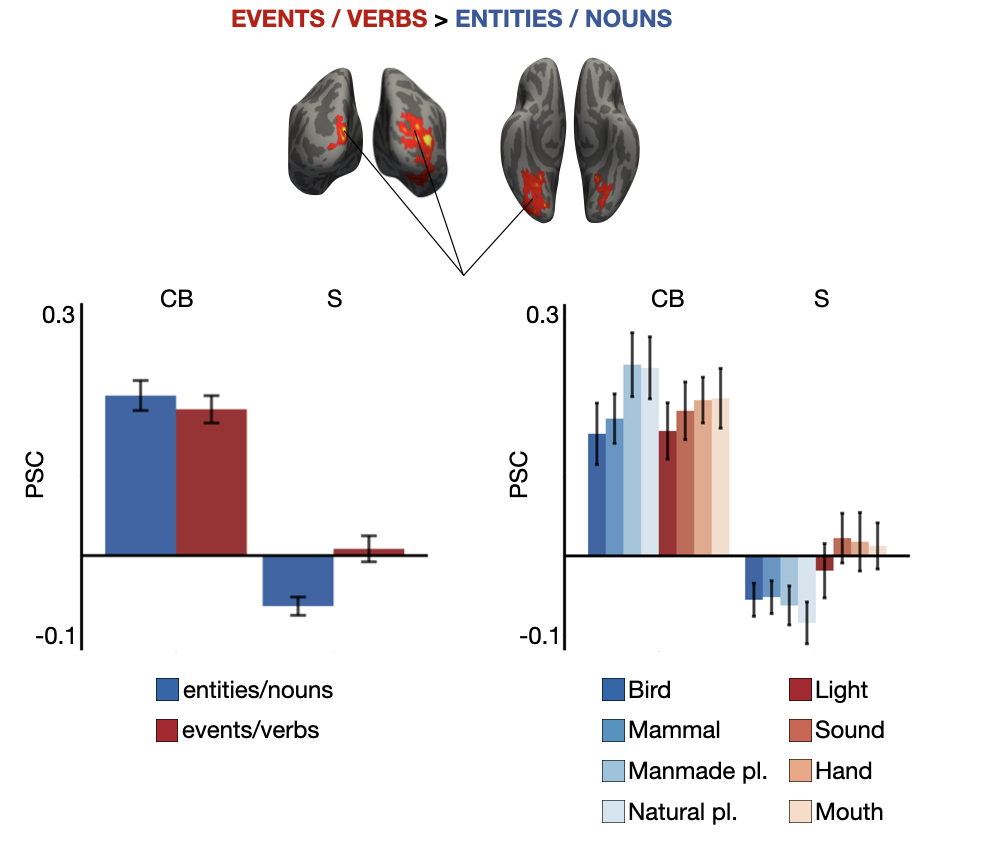

Supplementary Figure 8: Group-by-condition interactions of univariate verbs>nouns contrast in occipital cortices. Group maps are shown p<0.01 with FWER cluster-correction for multiple comparisons. Voxels are color coded on a scale from p=0.01 to p=0.00001. The left plot displays peak percent signal change averaged across all occipital regions in which group-by-grammatical class (events vs. entities) interactions were observed. The right plot displays peak percent signal change in these regions for each verb and noun category separately.
